# The view of the native gauges of blood pressure – focus on atrium (hydrodynamics and rheology)

**DOI:** 10.1101/037705

**Authors:** Yuri Kamnev

## Abstract

The precision of the inlet parameters depends on mechanism of response. Reflexes are satisfied with relative higher or lower but if the inlet information is presented by different parameters and the response is calculated according to some equation precision must reach the degree which does not slur over the results of calculation. At recent work the equation for controlling of circulation was suggested where the main inlet parameters are the arterial diastolic and vinous pressures and it becomes pertinent to analyze how organism can perceive these pressures with hydrodynamic accuracy. As far as the velocity pressure component of total pressure can not be detected by wall receptor of the rectilinear section of artery it was noticed that baroreceptors are located at the outer radius of the bend of central arteries and that is justified due to specific distribution of pressure. This phenomenon can be interpreted as a correction of measuring of static pressure with regard for velocity pressure component. Velocity pressure component of venous pressure is comparable with the one of arterial pressure but static components of venous and arterial pressures are incomparable and it is the fact that cannot be ignored when choosing the gauge. The possible method of measuring of pressure is based on observation that pressure-volume vector of the ventricular cycle is similar to a-loop vector of atrial cycle. Ventricular filling vector and x-trough vector show the behavior of viscous material but v-loop inserted into a-loop demonstrates typical viscoelasticity with creep. If viscous deformation of atrium at early relaxation possesses standard duration being stopped by transformation of viscous deformation into viscoelastic deformation the venous pressure can be measured in accordance with the value of viscous deformation. Measuring of pressure by viscous method implemented by atrium has the advantage comparing to measuring by baroreceptor with elastic sensor. Early relaxation of atrium which reveals coefficient of viscosity corresponds to ventricular relaxation and its coefficient of viscosity but the latter is liable to different biochemical shifts. Such shifts influences the atrial coefficient of viscosity either and the values of venous pressure measured by viscous method will be more accurate for calculations because coefficient of viscosity participates in the equation being not estimated in organism.

## INTRODUCTION

Measuring of blood pressure by organism itself is proved now by the existence of responses like reflexes that change the heart rate or alter production of some hormones. Sensitivity and methods of measurement must depend on the existing of native coordinator (computational program) that assesses information, analyzes the balance and gives the response. The coordinators may superimpose one another being of different phylogenetic age and complexity but, nevertheless, we are able to distinguish each evolutionary layer as something separate. Mathematical modeling is the unique reliable approach that afford us to see the integrity of bonds which are abstracted from the excess of many other bonds. Consequently, the task, for the first time, is to look for the facts and observations which are hardly explained till now but will gain almost the evidence after participation at modeling (nevertheless, the evidence appears only after the two further steps – the interference into the natural regulation and the construction of artificial analogue). Mathematical model, that was advanced recently, pretends to coordinate several circulatory parameters by the equation deduced; blood pressure (venous and arterial diastolic), presented in it, is invested in numerical form and the values introduced for calculations variate the duration of ventricular diastole [1,2]. Consequently, the data from pressure sensors must not be accepted in relative form – like “higher” or “lower” that is sufficient for reflexive responses ensuing. Hence, if we want to scrutinize the ways of achieving the numerical precision by native gauges of blood pressure, firstly, it is necessary to examine the object of measurement, – i.e. the structure of the flow in the vessel with regard for measuring of pressure, – secondly, it is necessary to imagine how the nature can excel in selection of methods for gauging due to peculiar characteristics of pressure and the points of application of the data.

The definition that can be favored is the following: Pressure is measured by converting the physical phenomenon to an intermediate form, such as displacement, which can be measured by a transducer. The only comment is to be added, however: all gauges of the direct measurement of pressure exploit elastic deformation (displacement) of sensor – but there are exist viscous and plastic deformations.

## METHODS

The following hydrodynamic and rheological approaches are necessary for giving explanation of the difference between measuring of pressure of the moving blood in arteries and veins, including not only the levels of energy of these two flows but also the purposes of information about pressure values.

**Fig. 1.**
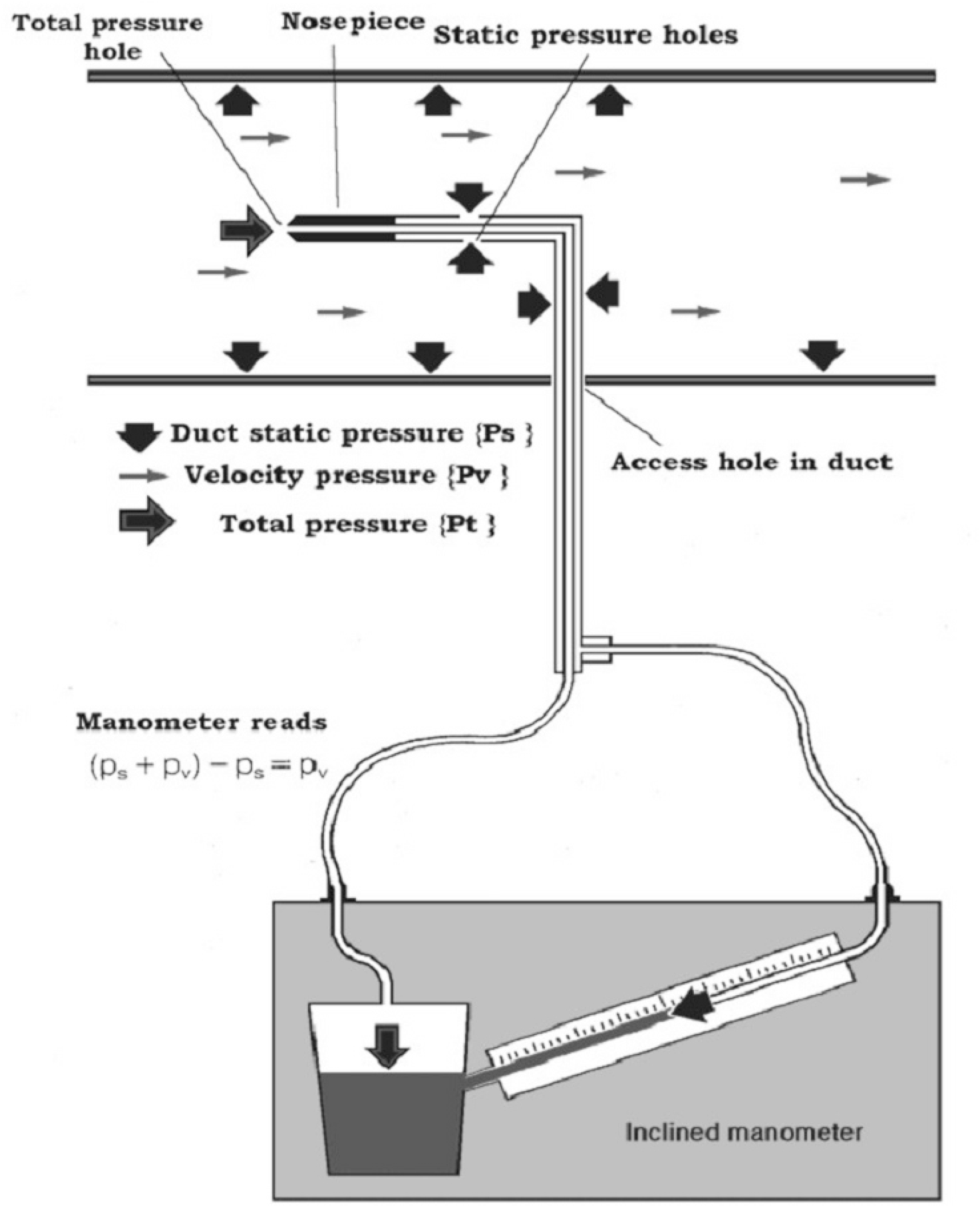
(extracted from [3]). Laboratory measurement of pressure in the duct where the liquid flows: vectorial presentation of the compound (total) pressure with regard for locations of the detectors.

1. **Static pressure and velocity pressure**. Fig.1 shows that the gauge inside the duct is subjected, firstly, to static pressure and, secondly, to additional dynamic pressure equal to the velocity pressure due to the impact; the algebraic sum of the static pressure and the velocity pressure is called the total pressure. A tube placed in a duct facing into the direction of the flow measures the total pressure; if frictional losses are neglected, the mean total pressure at any cross section throughout the duct system is constant. The determination of static pressure is carried out through the series of holes positioned at right angles to the flow (cited from [3], abridged).

**Fig. 2.**
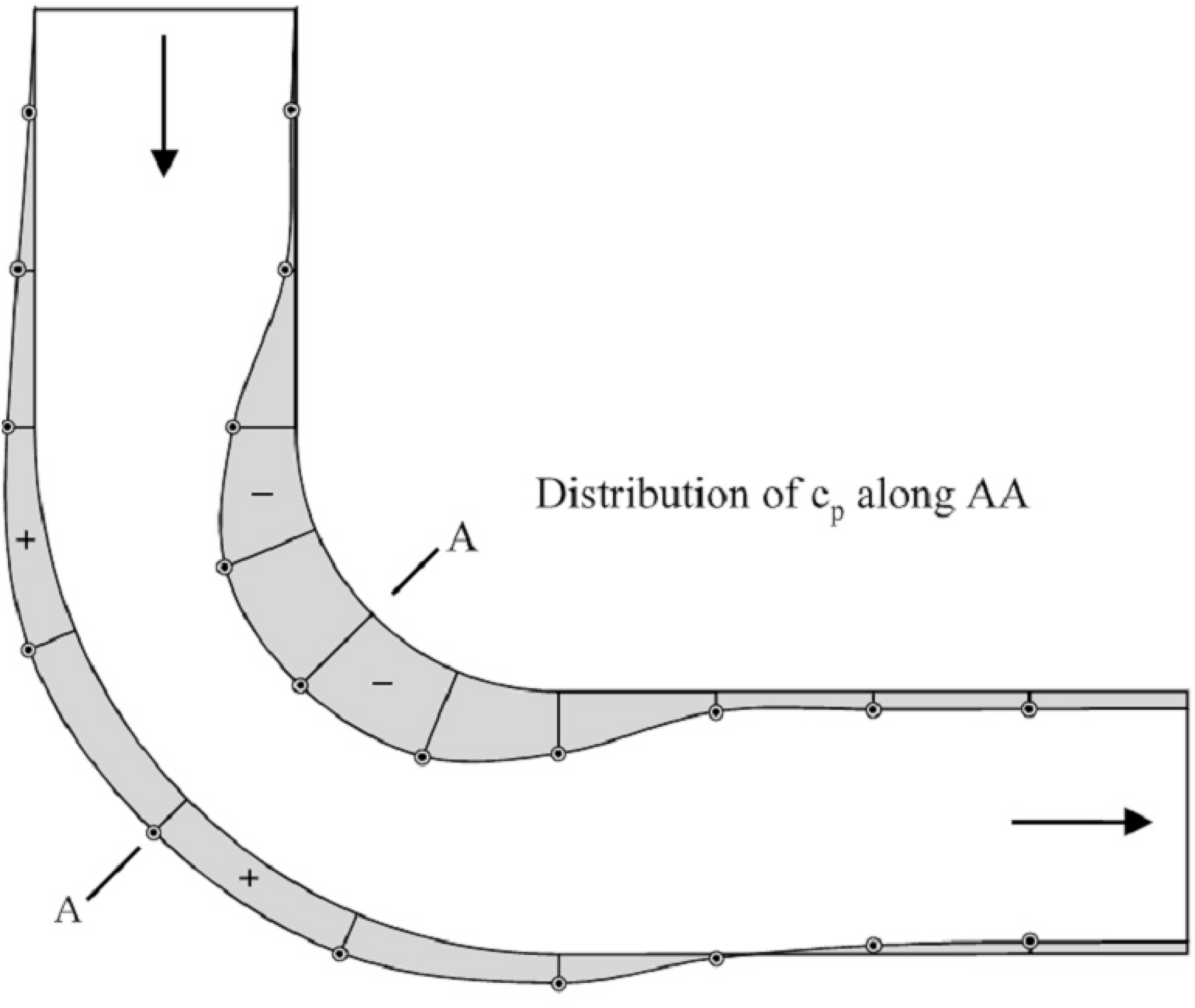
(extracted from [4]). Local elevation and drop of pressure near the walls of inner- and outer-radius of the duct with 90° bend (see text for details).

2. **Ducts with 90° bend**. Fig.2 plots the pressure coefficient Cp along the inner- and outer-radius walls. 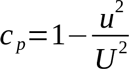, where U is the uniform velocity of the flow approaching the bend, and u is the streaming velocity at radius taken from the center of curvature of the bend [4]. The flow is accelerated in regions near the inner-radius wall due to the favorable longitudinal pressure gradient; conversely, the decelerated flow is observed in regions near the outer-radius wall because of the developed adverse pressure gradient. The outer bend is referred to as the pressure side and the inner bend as the suction side. The pressure obtained at the inner bend is seen to decrease gradually and monotonically; the dramatic pressure rise along the outer bend results in the aforementioned adverse pressure gradient. Such an adverse pressure gradient developed along the outer bend may cause the streamwise separation to occur (cited from [5], abridged).

**Fig. 3.**
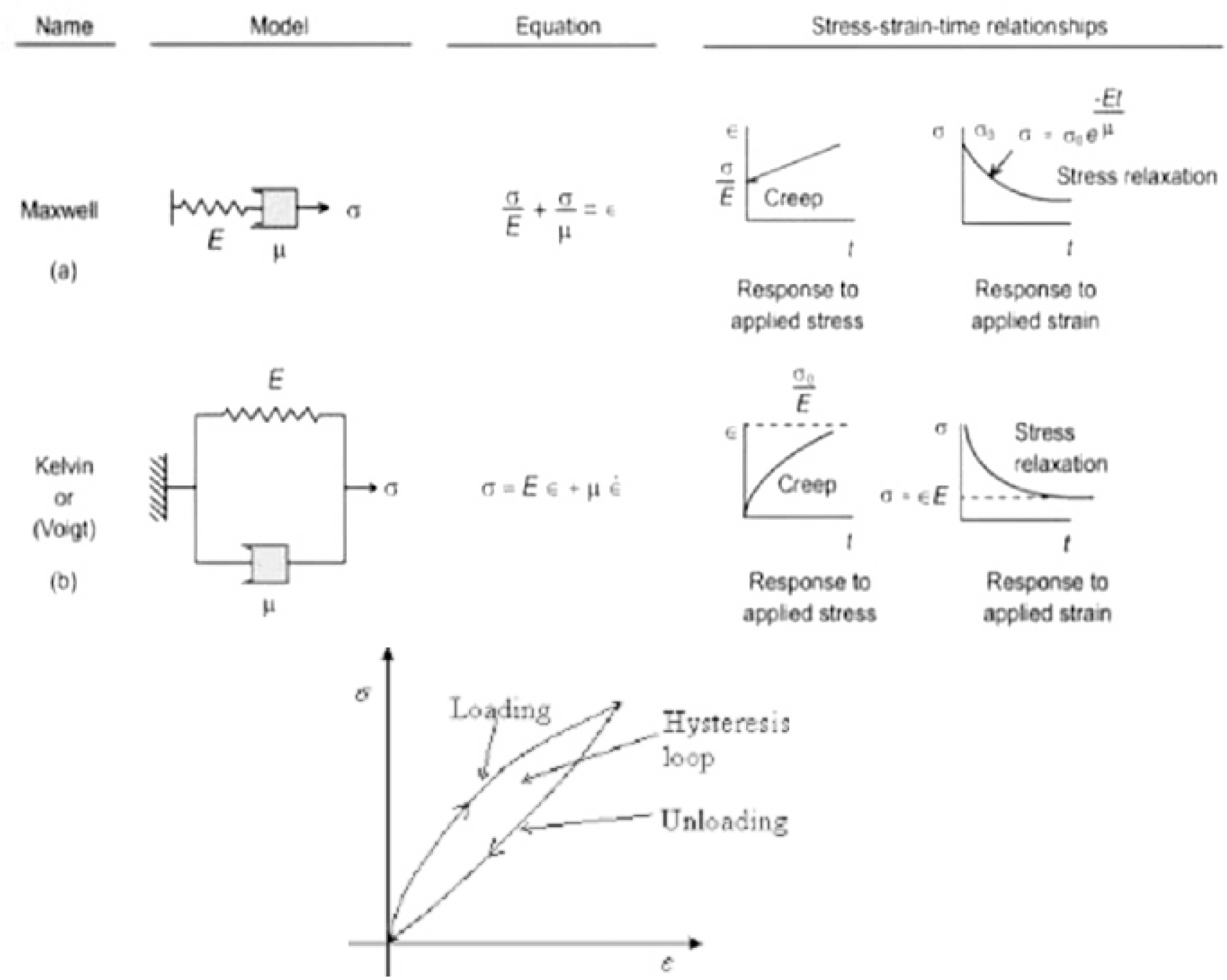
The upper part (extracted from [6]) presents the basic characteristics of complex materials which combine elasticity and viscosity. In both models elasticity acquires time-dependence; stress is no more an independent variable. The lower part pictures reversible reaction of loading-unloading with energy loss; the loop approximates the behavior of viscoelastic material.

3. **Viscoelasticity and creep**. The upper part of Fig.3 illustrates the appearance of time-dependance of stress and strain after the adjusting of viscous component to the elastic one – as far as viscous deformation is time-dependent but elastic one is not. Both Maxwell model and Kelvin-Voight model produce phenomenon of creep due to the work of viscous element and which is mostly responsible for energy loss that is being demonstrated by hysteresis loop. Nevertheless, the creep originated by Maxwell model can be interpreted as safety device because the growth of strain is linear while the developing of deformation; on the contrary, Kelvin-Voight model depicts the non-linear increasing of strain with steep growth of strain during early period of loading which may lead to rupture of material quickly. Let us compare both models of viscoelastic deformation with viscous deformation: viscous deformation is the function of two independent variables – stress and time; any viscoelastic deformation demonstrates the intermediate dependance – stress depends on time; consequently, the time is the only independent variable although deformation depends on time and on the value of stress either. Therefore, it is very difficult to control viscoelastic deformation by means of two factors of influence (stress and time); contrarily, viscous deformation is absolutely dirigible while controlling stress and time. The difference is very important if you want to regulate the filling of the volume.

The lower part of Fig.3 shows the abstract hysteresis loop of viscoelasticity but the real configurations may vary; however, vector of loading always occupies the superior curve of the loop and it is directed clockwise; vector of unloading is also directed clockwise (along the inferior curve). In other words, the deforming is a respectively high-energetic process which needs the indraft of energy for storage and, vice versa, the recovery of the essential shape is the process which spends (and partly dissipates) energy stored.

**Fig. 4.**
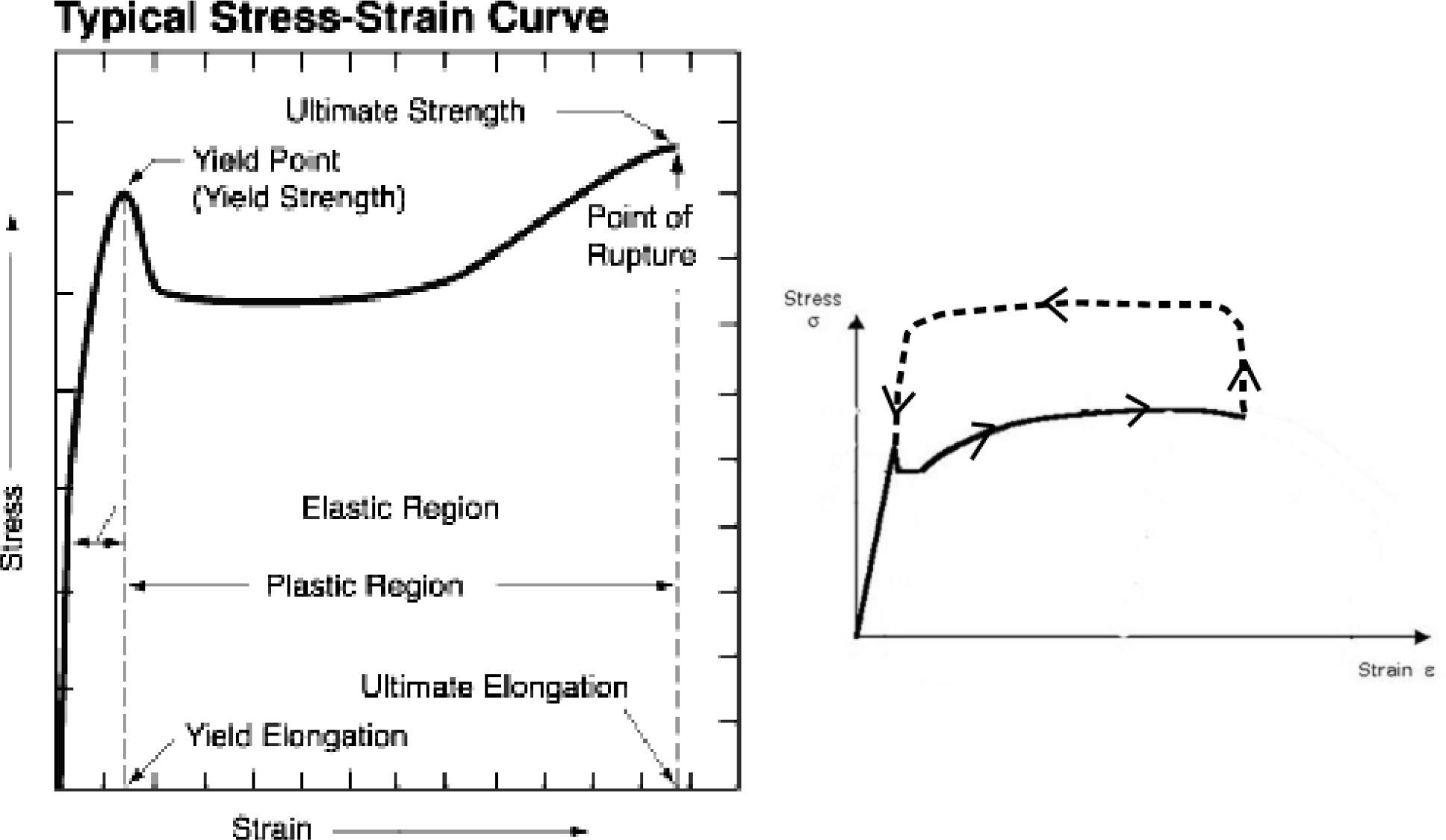
The left diagram describes the elastic-plastic boundary experiment; the horizontal section is the description of viscous flow while the plasticity itself (permanent deformation) appears only after the withdrawal of yield strength. The right diagram partly copies the graph from the left diagram indicating the development of process by arrows; the route (the dotted line with arrows) of hypothetical restoration of initial form of object is added; vertical rise is the imaginary fragmentation of material at higher stress, horizontal directed to the left is the subsequent forming of the shape that has preceded the yield-point strain; consequently, the withdrawal of yield strength (vertical drop) will not result in plastic deformation.

4. **Flow of solids**. Trough-like section of the curve (plastic region) at Fig.4, left part, is described by the system of constitutive equations which include the description of the flow of the viscous fluid and conditions concerning the yield limit is τ_s_. Therefore, if the reversible deformation passed over yield point the energy is spending now to molecular reconstruction and the process can hardly be called “storing of energy” – the yield only absorbs energy and viscous deformation is developing while stress slightly decreases, keeps on the steady state or increases (within limitations). As far as deforming does not stop (while yield level of energy is maintained) the energy of stress is continually reconstructing the material and any attempt to reduce stress below the “bottom of the trough” does not result in recovery of the form: the energy was not stored but it was wasted. Therefore, the recovery of initial form can not be reached by unloading because the above mentioned reconstruction (plastic change of initial form) determines the new lowest starting point – the new shape of object – for the future elastic deformation. By the way, if we reduce, or even neglect, the elastic region (Fig.4), – which is energetic staircase to the trough of the yield, – and which depends only on the nature of the material, – we may find out that the process of flowing itself (viscous deformation) is ultimately economical: viscous deformation does not need the gradual elevation of stress (like elastic or viscoelastic deformation). Certainly, the elevation of stress will accelerate viscous deformation bringing it quicker to the breaking point, – and the absorbed energy is impossible to take back, – but if we want to imitate the flow of solids by means of some device the deformation can be achieved due to ultimate economy of energy.

The question is: if it is impossible to restore the initial form by the diminishing of stress – may be it is possible to restore it by the indraft of energy? It may not be stress but it may be temperature – imagine that you fuse the overstretched spring and mould a new one. Or – if without rising a temperature – we may imagine increasing of stress (Fig.4, right part, dotted line) which rupture the flowing material down to structural elements; these elements will be undergone reconstruction aimed to liquidation of viscous deformation – the dotted line returns to the yield point. Certainly, such process needs either the indraft of energy or the specific system of, firstly, fragmentation of the fluid-like material at the molecular level (while stress increasing) and, secondly, reconstruction of the shape due to reverse direction of flow (while high stress persists) and, thirdly, the restoration of solid structure (while stress decreases).

## RESULTS

The most important baroreceptors of systemic circulation are located in carotid sinus (at the bifurcation of external and internal carotids) and in the aortic arch; both zones occupy the bends of vessels and there is no information about the reception of systemic arterial pressure from rectilinear sections of big arteries. Arterial baroreceptors of pulmonary loop are located in the vicinity of the main bifurcation of the pulmonary artery, or in the right and left branches between the main bifurcation and the origins of the lobar branches; no receptors have so far been found in the pulmonary trunk proximal to the main bifurcation [7]. The above trivial data correspond to hardly trivial supposition that controlling of circulation is in need not only of values of static pressure. The latter one can be measured at the wall of rectilinear section of artery but measurement of the velocity pressure requires some probe (catheter) inside the vessel (see Fig.1) – and it is not feasible in nature. The solution, as the first approximation, may be offered the following way: pressure sensors must be places on the outer-radius wall of anatomical bends of arteries. Total pressure can be registered here, with maximal attainable accuracy, by the wall sensors – due to revealing of velocity pressure component directed against the outer-radius wall.

Low-pressure receptor zones are located in the venae cavae, pulmonary veins and in the atria (in the right atrium – at the junction of venae cavae, in the left atrium – at junctions with pulmonary veins). Venous pressure, in the vicinity of the heart, is 2-5 mmHg (or even negative when we inhale but values below atmospheric zero obviously relate to the measured static component of venous pressure) and it is approximately 15-fold lower comparing to diastolic arterial pressure; velocity flow rate in venae cavae is no faster than 20 cm/sec and velocity flow rate in great arteries is maximally 30-50 cm/sec. Therefore, velocities are comparable but pressures are not. It is obvious that the portion of velocity pressure in total pressure of the venous flow is much higher then the portion of velocity pressure in total pressure of the arterial flow. Certainly, it needs accurate experimental assessment but a statement that measuring of pressure by wall sensors at the bend sections of arteries raises the precisiness (due to detection of velocity pressure component) is well-reasoned. Nevertheless, we are speaking about the correction of measurement of static pressure – contrarily to almost dominant portion of velocity pressure in total pressure of the venous flow (the statement needs accurate experimental assessment either). As for theoretical comparison – if the portion of velocity pressure prevails then the methods which are good for correction are inappropriate for evaluation; although stretch receptors at junctions of central veins and atria are really activating reflexes of Bainbridge’s type. In some cases, as we know, the central venous pressure, measured via catheter, may raise very high – and in that cases we are speaking about total pressure with static component significantly increased, measured via catheter. Such reflex causing heart rate acceleration can be considered some safety device (and very early phylogenetic reflex) which protects the ventricle – relaxing like viscous material, as we assume – from the rupture caused by immediate shortening of diastole. The same action (regulating of the duration of diastole) – but as a precise response on an accurate measuring of venous pressure when the velocity pressure component prevails, – is expected in the form of a system which combines the process of measuring of venous velocity pressure and process of relaxation of ventricle.

**Fig. 5.**
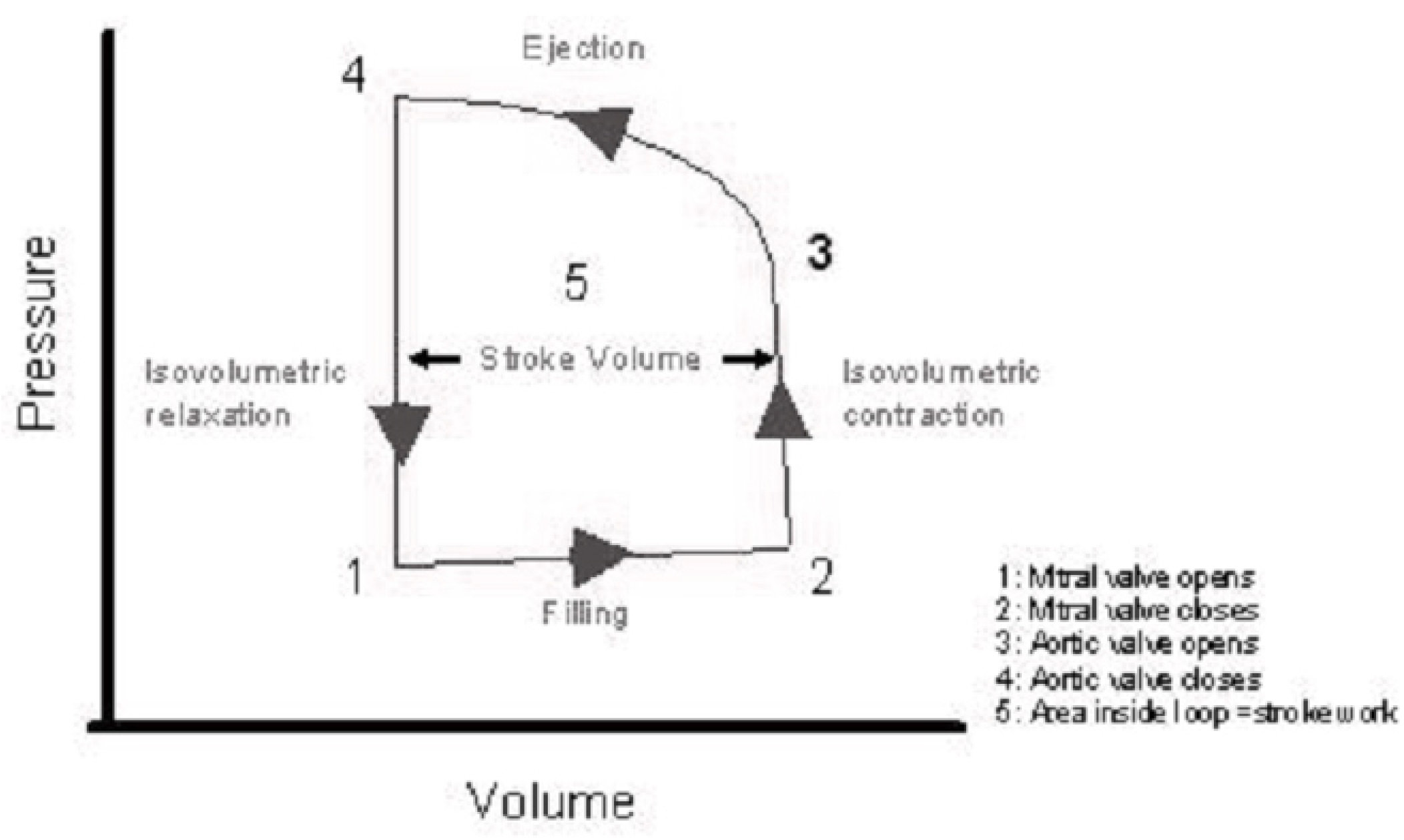
Schematic pressure-volume loop of the ventricular cycle. Counterclockwise direction of vector indicates that the restoration of the initial shape of the ventricle (the shape before filling, i.e. before viscous deformation) accomplishes with the indraft of energy, i.e. elevation of pressure (stress) while muscle contracts. Raising level of stress (isovolumetric contraction) damages the previous structure of dilated ventricle (right vertical) and then the shortening of the muscle reconstructs the molecular structure and shape of ventricle before filling (upper horizontal); cycle is ending with the subsequent drop of stress, i.e. isovolumetric relaxation (left vertical).

**The coincidence of loops of σε-vectors**. Hypothetical redirection of viscous process (flow of solids) from plastic hardening to some material structure modifying (which restores the initial form of an object bringing σε-vector back to the yield point) is plotted at Fig.4, right part, by dotted line with arrows. Such process needs the increasing of stress which, however, must not break the material in terms of rupture, – and we do not know such examples in classical rheology. Let us have a look at Fig.5 which depicts the well-known pressure-volume (= σε) loop of ventricular relaxation-contraction cycle. The lower horizontal vector directed counterclockwise can be interpreted only as viscous deformation (but not viscoelastic creep which is shown at Fig.3 and which is similar to the atrial v-loop at Fig.6). The rest arc vector directed counterclockwise demonstrate that the above mentioned hypothetical process was realized in ventricular muscle contraction and isovolumetric relaxation of ventricle.

As far as viscoelastic process was mentioned already it is pertinent now to turn to pressure-volume atrial cycle (Fig.6) and repeat that v-loop is nothing but hysteresis loop of viscoelastic deformation. The matter of interest and profound analysis is a-loop and its junction with viscoelastic loop. Imagine v-loop is ultimately diminished (down to disappearance); in that case, a-loop gains similarity to ventricular loop and we may interpret confidently the horizontal counterclockwise vector as viscous deforming of early atrial relaxation. Respectively, – when v-loop is absent, – viscous deformation immediately turns to muscle contraction (isovolumetric contraction at first) as it is in case of ventricular cycle. In ventricle the transition from relaxation to contraction is stipulated by impulse of excitation (which regulates the duration of diastole), but in atrium, – if v-loop is absent, – no special impulse appears. Consequently, v-loop originates from the necessity of waiting to further excitation – from the appearance of some break (pause) between atrial viscous relaxation and atrial contraction – under conditions of continuous stress of venous pressure which will inevitably rupture viscous material. Therefore, some biochemical operator must exist that converts viscous process of relaxation into viscoelastic one – which adds protective characteristics to stand the stress. Switching on of such biochemical operator can be connected, quite probably, with previous impulse of excitation and further contraction of atrium as far as no other event controls the beginning of viscous deformation. Since viscous deforming is the function of two independent variables (stress and time) very interesting engineering device suggests itself: if the time-interval from the beginning of atrial relaxation (or even from sinus impulse) and up to the moment of conversion of viscous relaxation into viscoelastic relaxation is fixed – it is quite possible to measure stress by means of gauging the level of filling (degree of dilatation) of atrium at the end of the period of viscous relaxation. It would be the original method of measuring of pressure by means of viscous deformation but not by elastic deformation of membrane (sensor).

**Fig. 6.**
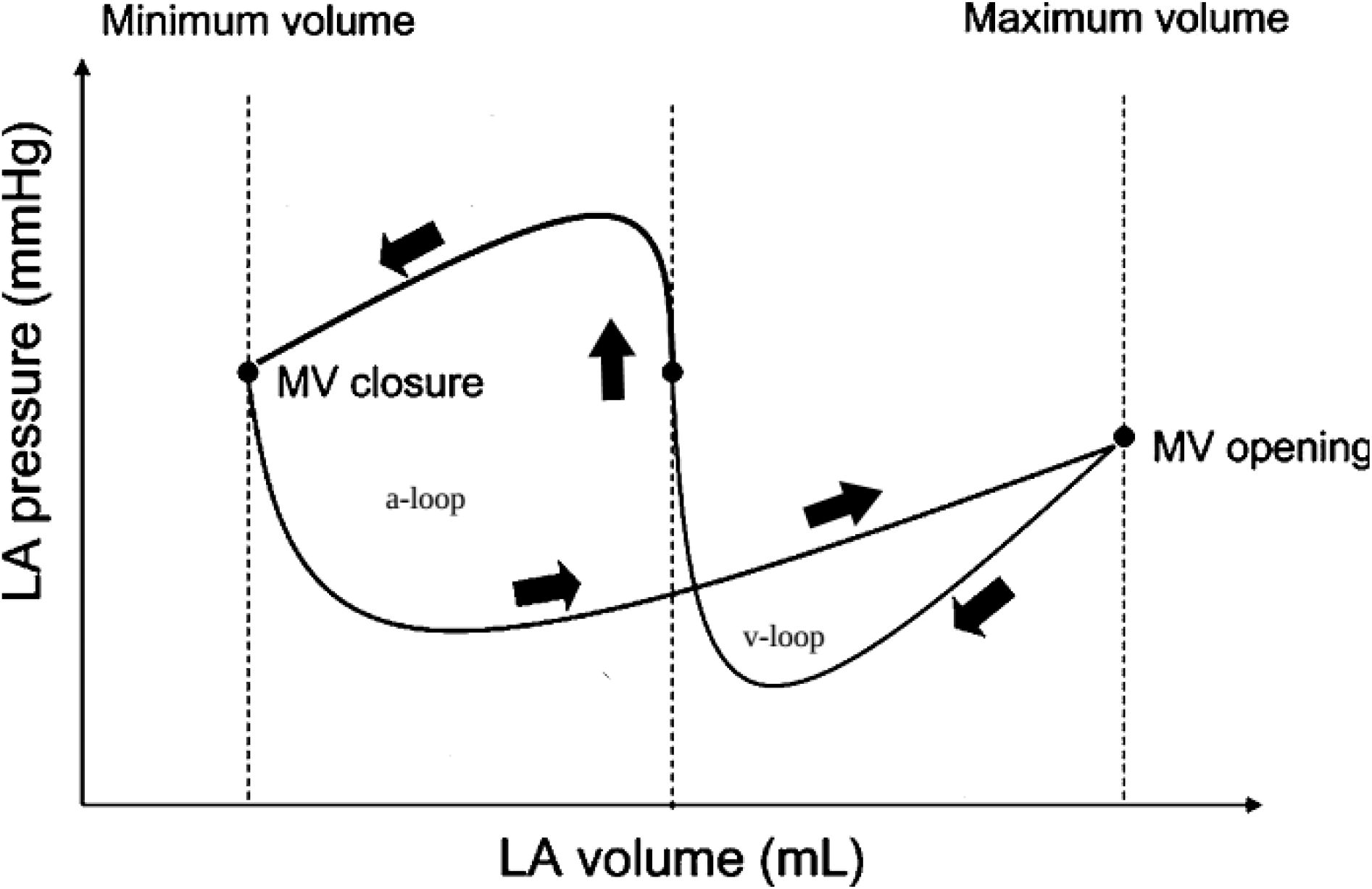
Schematic pressure-volume loops of the atrial cycle (left atrium). A-loop is developing according to the counterclockwise direction of vector and it resembles to the ventricular pressure-volume loop; v-loop is developing according to clockwise direction of vector and it resembles to viscoelastic hysteresis loop.

**Synchronization of atrio-ventricular valve closure and the beginning of atrial relaxation**. The possibility of measuring of venous pressure by means of viscous deforming process (that is principally another method comparing to measuring of pressure by means of elastic displacement functioning at all pressure sensors) puts the question: what is the phase of cardiac cycle that is optimal for measuring. In other words, what period of atrial cycle the total venous pressure, – with the dominant of velocity pressure component, – does influence the wall of the vessel (of the atrium) similar to the influence on the ventricular wall? It may seem that we have changed the criterion and does not look for the conditions to register the total pressure completely as a sum of static and velocity pressures. However, the ventricle – as the ending of venous pipe – trivially can be considered even the better receptor of velocity pressure component then the bend of the vessel (the direction of the vector of velocity flow rate is orthogonal to the imaginary plane of the vessel ending). Therefore, the answer to the question is the following: the better phase is when the atrium behaves like ventricle itself and, consequantly, such phase is situated just after the atrial contraction. At this period venous flow is depraved from transition to the ventricle because valve is closed, – the event which needs synchronization between two systoles (atrial and ventricular). Under this condition the atrium, at early relaxation, will be influenced by the same stress as the ventricle during the ventricular diastole. As it was mentioned, viscous deformation of atrium can obviously be undergone to the stress of venous pressure very shortly; standard time interval of viscous deformation (organized by “the border” between viscous and viscoelastic deformation) results in possibility of measuring of venous pressure. Consequently, synchronization of atrial and ventricular contractions and relaxations is needed; hence, fixed atrio-ventricular delay of impulse gains very rational explanation.

**The emergence of viscoelasticity after viscosity during atrial relaxation**. Measuring of pressure by means of viscous deformation claims standard duration of viscous deforming; firstly, the inception of this period is caused by the completion of atrial contraction and, secondly, the inception of this period is synchronized with atrio-ventricular valve closure. The whole period of viscous deforming corresponds to x-trough section at pressure-volume diagram (Fig.6) and, respectively, the end of this period is marked by the beginning of viscoelastic loading. The change of material behavior is proved by subsequent viscoelastic unloading which turns vector clockwise – i.e. the material demonstrates the recovery of shape and returning of the stored energy (v-loop). As a matter of fact we do not know what trigger does switch the imitation of viscous deforming over to imitation of viscoelastic one during atrial diastole. Nevertheless, we may interpret v-loop as an insertion of viscoelastic behavior of material (v-loop, clockwise direction of vector) into another mode of deformation-and-restoration (a-loop, counterclockwise direction of vector) which is, so to say, a representative of ventricular mode of deformation-and-restoration (vector of ventricular cycle is directed counterclockwise). It is ought to remember that specific process of liquidation of plastic deformation by means of muscle contraction is biological process without analogues in classical rheology. Thus, two goals are achieved by the above mentioned insertion of v-loop into a-loop: firstly, the standard duration of x-trough appears despite the wide range of duration of atrial diastole (which corresponds to the range of ventricular diastole, i.e. to diapason of the heart rate, respectively); secondly, viscoelastic deformation plays the protective role for the rest part of atrial diastole whereas x-trough (viscous pressure sensor) can be influenced by stress of total venous pressure very shortly.

**Equal liability of ventricular relaxation and early atrial relaxation to different factors**. Since we assume that relaxation of ventricle and early relaxation of atrium is the viscous process it is pertinent to refer the knowledge about ventricular relaxation, undergone to different factors, to the early atrial relaxation either. From rheological point of view the interference in such relaxation is the change of viscosity of material (change of imitation of viscous behavior of material). It is not worth enumerating biochemical shifts that can alter the relaxation [8]; more important is to comprehend that the influence on relaxing ventricle and the influence on the early atrial diastole must be identical. Simple formula of viscous deformation is the following 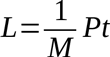, where L is relative viscous deformation, M is the coefficient of dynamical viscosity, P is pressure and t is time. (The following example presupposes the acquaintance with our previous works [1,2].) Imagine the venous pressure was measured by some stretch-sensitive receptor and, in one case, the same value was analyzed in cooperation with M = a and, in other case, was analyzed in cooperation with M = b; the information about the difference can not be taken into consideration because only pressure values (and duration of diastole) are involved in regulation. Thus, signals from stretch-sensitive receptors indicate the steady venous pressure – but the end-diastolic volume is changed, systolic pressure is changed and no correction concerning the new duration of diastole is calculated; consequently, the real venous pressure (residual pressure) is changed and stretch-sensitive receptors will register the alteration of venous pressure – although initially venous pressure was not changed (but only M); now the correction will happen and circulation will be found on some other level of circulatory pressures. Contrarily, if vinous pressure was measured by viscous receptor we gain the new information – as if venous pressure has been changed (whereas M was changed but not the pressure) and this information gives the command to change the duration of diastole (in order to restore equipoise as far as no information about some change of arterial pressure has come – which may justify the change of ventricular filling); as a result – the end-diastolic volume will not be changed due to the correction of duration of diastole, systolic pressure will not be changed and venous pressure (residual pressure) will not be changed; viscous pressure receptor again assesses it in ensemble with altered M and detects the value identical to the value from the previous cycle, i.e. as changed but equal to the previous one (again the real pressure is not changed but only M); nevertheless, the correction of the heart rate was carried out already and, therefore, the level of circulatory pressures remains the initial.

Hence, the observation follows that seems contradict with the premise expressed about the obligatory precision when the data is applied to calculations, – the precision alleged to be due to absolute values at least. It was true when we deal with arterial pressure measurement which is accomplished by receptors with elastic deformation of sensor, – i.e. we suppose that elastic properties (elastic module) of sensor are invariable. But the above example, which is based on hypothetical measurement of venous pressure by means of viscous deformation of atrium, suggests viscosity as the working property of material of sensor. As far as viscous deformation is associated with early relaxation of atrium (behaving like relaxing ventricle) such viscous sensor will be prone to many biochemical shifts and, consequently, may change its viscous properties. Moreover, there is no feedback concerning any changes of relaxation; consequently, the information about viscous process (as a pressure gauge) must be taken into consideration in pair with coefficient of viscosity (= relaxing properties) which alters without any quick signals. It turns out, – and it goes from the above example, – that the relative values of venous pressure are more correct for the final calculations then the absolute values gained, for instance, from stretch-sensitive receptors. Therefore, in one case (arterial pressure), the precision is achieved by more accurate measuring of absolute values, and in other case (venous pressure), – when measuring of pressure depends on variability of viscous properties of atrial wall (deterioration of accuracy), – “the noise” produced by fluctuations of viscosity appears to be the common factor for ventricular relaxation either – and thus final calculation does not suffer.

## REFERENCES

1. Kamnev Y. 2015. Viscous deformation of relaxing ventricle and pulsatile blood propelling (numerical model). bioRxiv doi: 10.1101/009522

2. Kamnev Y. 2015. The conversion of systolic volume into systolic pressure (the development of numerical model). bioRxiv doi: 10.1101/011536

3. Matthews G.J. How to measure pressure. How to measure flow. Airflow Developments Limited. www.tsi.com

4. Markland E. A first course in airflow. 1976 (2011), ch.8. Flow around a bend in a duct. web.iyte.edu.tr

5. Sheu W.H.Tony, Tsai S.F. Vortical flow topology in a curved duct with 90 bend. Proceedings of the 4th WSEAS International Conference on Fluid Mechanics and Aerodynamics, Elounda, Greece, August 21-23, 2006 (pp121–129)

6. Ravindra Gettu. Module-3, lecture-10. Rheology of liquids and solids. textofvideo.nptel.iitm.ac.in/…/lec19 pdf

7. Coleridge J.C.G., Kidd C. Electrophysiological evidence of baroreceptors in the pulmonary artery of the dog. J.Physiol. (1960), 150, pp.319–331

8. Chemla D., Coirault C., Hebert J.-L., Lecarpentier Y. Mechanics of relaxation of the human heart. News Physiol.Sci., v.15, April 2000, pp.78–83.

